# Conserving Biodiversity in Coffee Agroecosystems: Insights from a Herpetofauna Study in the Colombian Andes with Sustainable Management Proposal

**DOI:** 10.1101/2023.03.07.531390

**Authors:** Juan Camilo Ríos-Orjuela, Nelson Falcón-Espitia, Alejandra Arias-Escobar, Dennys Plazas-Cardona

## Abstract

Amphibians and reptiles are important indicators of ecosystem health, yet their populations are declining worldwide due to habitat loss and climate change. Agroecosystems, such as coffee plantations, can provide important habitat for these species. We conducted field surveys in the Sumapaz region of Colombia to identify the microclimatic variables that influence the diversity and abundance of herpetofauna in coffee crops. The canonical correspondence analysis revealed that leaf litter abundance, crop area, and age category were the most important structural variables for determining herpetofauna diversity. Our findings suggest that coffee plantations function similarly to secondary vegetation, and maintaining a thick layer of leaf litter is critical for establishing complex and structured animal communities. This study proposes a set of sustainable agricultural management principles to promote the existence of amphibians and reptiles in coffee crops. By adopting these practices, it is possible to prevent the decline in the population of amphibians and reptiles due to the expansion of the agricultural frontier, as seen in other coffee-growing regions. The findings of this study contribute to a better understanding of how to balance agricultural production and biodiversity conservation in the context of agroecosystems.

* *Una versión en español de este documento puede ser encontrada como material suplementario* / a Spanish version of this paper can be found as supplementary material.

## INTRODUCTION

Land use change for agriculture is one of the main drivers of biodiversity loss in the world (Manson et al., 2008). As coffee is an economically important crop in world agriculture, the impact of its cultivation on the ecology and diversity of certain animal groups has been investigated in altered environments (Philpott et al., 2008; Şekercioglu et al., 2019). Studies have shown that environmental-specific characteristics and adjacent ecosystems influence changes in biodiversity (Cisneros et al., 2016; Newbold et al., 2015). Regarding herpetofauna, research into the effect of different crops on species turnover and local extinction risk in Latin America has focused on amphibians for the most part (Murrieta-Galindo et al., 2013; Santos-Barrera and Urbina-Cardona, 2011); with limited reports on reptiles (Mendenhall et al., 2014). Information gaps exist in the neotropics, particularly in Colombia, a country with high amphibian and reptile diversity, a broad coffee production culture, and a lack of studies on the impact of coffee cultivation on herpetofauna.

The Cerro Quininí Protected Forest Reserve (RFPCQ) is a vital conservation area in the Sumapaz province (Cundinamarca, Colombia) and forms part of the crucial Chingaza-Sumapaz-Guerrero corridor in the Eastern Cordillera of the Andes (Sguerra et al., 2011). Despite being a protected area, approximately 90% of the associated land is privately owned and has been utilized for cultivation and grazing, with coffee cultivation occupying around 80 ha (Alcaldía Tibacuy, 2016). As such, new conservation and development methods have been explored in agroecosystems like the RFPCQ, as they hold a diverse range of species and have been historically overlooked by conservation efforts (Rojas Sánchez et al., 2012). Given its environmental conditions, the RFPCQ holds the potential for high species diversity, but habitat loss impact has not been evaluated at the ecosystem or species level. Available data suggest 29 amphibian and 88 reptile species could be present, including 11 threatened amphibians and three reptiles (IUCN, 2021; Morales-Betancourt et al., 2015; Rueda-Almonacid et al., 2004). Thus, further research and conservation efforts are needed to protect the RFPCQ’s diverse array of species.

To consider biological diversity in the middle altitude Andes range, the coffee agroecosystem must be considered. Despite the alterations to the original ecosystems, some ecological services are still maintained in coffee crops, allowing different organisms to persist in these areas. Agricultural intensification is a trend in Latin America, with 48% of Colombian coffee-growing municipalities experiencing production changes due to replacing shaded crops with unshaded crops (Guhl, 2006; Rice, 1999; Rojas Sánchez et al., 2012).

Shaded coffee plantations in RFPCQ may serve as connection zones between less intervened ecosystems since they help maintain complex floristic and ecological structures, harboring wildlife species (Manson et al., 2008; Moorhead et al., 2010; Pineda et al., 2005), and even showing comparable diversities between shaded coffee plantations and natural forests (Tejeda-Cruz and Sutherland, 2004). However, limited research exists on herpetofauna diversity in agroecosystems in Latin America and Colombia (Hoyos-Hoyos et al., 2012; Moguel and Toledo, 1999; Murrieta-Galindo et al., 2013; Pineda et al., 2005). Therefore, to mitigate the negative impact of human activity on local wildlife and preserve the Andean ecosystems, it is essential to document the diversity and interaction of amphibians and reptiles with the local production systems.

Our objective was to assess the impact of coffee crops on the herpetofauna diversity in the RFPCQ. To achieve this, we had three main goals: (1) document the herpetofauna diversity in different vegetation covers associated with coffee production in the RFPCQ; (2) evaluate the relationship between the structural characteristics of coffee plantations and the amphibian and reptile community composition; and (3) propose a management strategy for coffee production systems that supports the presence of herpetofauna.

## METHODS

### Study area

The Cerro Quininí Protected Forest Reserve (RFPCQ) is in the municipalities of Nilo, Tibacuy, and Viotá (Cundinamarca, Colombia), between 1050 and 2133 m.a.s.l in the Eastern Andes (Castellanos-Menjura et al. 2019). The region has a temperate climate, with an average temperature of 19.2°C and a bimodal rainfall pattern (Corporación Autonoma Regional de Cundinamarca, 2016). According to Holdridge’s classification (1947), the RFPCQ is in the low-montane rainforest and low-montane dry forest life zones to a lesser extent. With an approximate area of 1947 ha, it is one of the largest conservation areas in Cundinamarca (Sguerra et al., 2011).

Despite being a protected area, most lands associated with the RFPCQ (about 90%) are privately owned and used for mixed crops and grazing(Alcaldía Tibacuy, 2016; Castellanos, 2015; Corporación Autonoma Regional de Cundinamarca, 2016; Corporación Autonoma Regional de Cundinamarca and Andina., 2013). In 1987, the area was declared a Protective Forest Reserve to conserve natural resources and the environment, particularly important water sources at the request of the inhabitants (Ministerio de Agricultura de Colombia, 1987).

### Data collection

To cover the rainy and dry seasons in the region, a total of six field trips (three effective sampling days each) were conducted between June and December 2021. Sampling localities were established beforehand based on the natural, semi-natural, and transformed plant units in the area, as well as the coffee crops in the RFPCQ. The sampling included grasslands, coffee crops, secondary vegetation, and forest ecosystems (Table *1*; a complete description of each vegetation unit can be found in Box *1*). Two of the sampling points were outside the RFPCQ, in the villages of Pueblo Nuevo and Buenos Aires (Viotá municipality), selected due to the absence of conserved forest cover in the reserve, while keeping equivalence with the slope and the altitudinal range in which the RFPCQ is located. As well, cropping systems were characterized according to proximity to water bodies, frequency of maintenance, use of agrochemicals, total area, adjacent vegetation covers, presence of microhabitats such as leaf litter and rocks, and percentage of shading (Table *2*).

**Table 1.** Alpha diversity indices for vegetation cover units. This table presents the alpha diversity indices used to estimate the diversity of vegetation cover units. The first line states the number of polygons and the total area of the analyzed vegetation units. The value highlighted in bold indicates the highest diversity unit following Shannon’s index.

| Index | Open forest | Secondary vegetation | Wooded pastures | Weeded pastures | Clean pastures | Coffee crops | Urban areas |
| --- | --- | --- | --- | --- | --- | --- | --- |
| Number of polygons / Total Area | 3 polygons / 68.12 ha | 7 Polygons / 90.72 ha | 6 polygons / 39.16 ha | 1 polygon / 0.49 ha | 1 polygon / 0.8 ha | 6 polygons / 5.97 ha | 1 polygon / 0.07 ha |
| Taxa_S | 9 | 23 | 8 | 2 | 3 | 9 | 1 |
| Individuals | 20 | 55 | 15 | 2 | 5 | 36 | 1 |
| Dominance_D | 0.245 | 0.084 | 0.164 | 0.5 | 0.44 | 0.399 | 1 |
| Simpson_1-D | 0.755 | 0.915 | 0.835 | 0.5 | 0.56 | 0.600 | 0 |
| Shannon_H | 1.799 | **2.776** | 1.934 | 0.693 | 0.950 | 1.41 | 0 |
| Evenness_e^H/S | 0.671 | 0.698 | 0.864 | 1 | 0.862 | 0.455 | 1 |
| Brillouin | 1.373 | 2.307 | 1.436 | 0.346 | 0.599 | 1.155 | 0 |
| Margalef | 2.67 | 5.49 | 2.585 | 1.443 | 1.243 | 2.232 | 0 |
| Equitability_J | 0.818 | 0.885 | 0.93 | 1 | 0.865 | 0.641 | 0 |

#### Box 1. Vegetation cover types and their characteristics in the study area.

Summary of the vegetation cover types and their characteristics in the study area. The box includes the vegetation unit’s definition based on (IDEAM, 2010), the total area sampled, and the number of polygons sampled in each vegetation type.

**Open Forest:** can be defined as a vegetation community dominated by regularly spaced trees forming a dispersed canopy, with a height greater than five meters. This type of forest cover has either not been disturbed or has undergone selective intervention, preserving its original structure and functional characteristics. In this study, three open forest polygons with a total area of 68.12 ha were sampled in the villages of Buenos Aires (Nilo), Atalá (Viotá), and La Vuelta (Tibacuy). The Buenos Aires and Atalá polygons are larger and correspond to continuous forests, while the La Vuelta polygon is surrounded by secondary vegetation.

**Secondary Vegetation:** this vegetation type is a result of the natural succession process that occurs after the intervention or destruction of primary vegetation. It can be found in areas cleared for different uses, in abandoned agricultural areas, and in areas where natural events have destroyed the natural vegetation. Within the study area, this unit occupies a total area of 90.72 ha, distributed in seven polygons located in the villages of La Vuelta, Bateas, Albania, Capotes, and El Cairo (Tibacuy). The cover in these areas is characterized by mainly shrub and herbaceous vegetation with an irregular canopy and occasional presence of trees and creepers. These areas are usually surrounded by other coverages such as pastures and coffee plantations.

**Clean pastures:** these are defined as lands covered mostly by dense grasses such as those of the Poaceae family and are used for permanent grazing for two or more years. In our study area, we conducted sampling in a single 0.8 ha polygon located on the La Vuelta trail in Tibacuy, which is used for livestock maintenance.

**Wooded pastures:** these are areas of pastureland with trees irregularly distributed over the land and forming 30% to 50% of the total area. These areas were selected for sampling in six polygons covering a total area of 39.16 hectares, located in Buenos Aires (Nilo), Capotes, La Vuelta, and Bateas (Tibacuy).

**Weeded pastures:** these are areas where grasses and weeds form associations of secondary vegetation, which are typically less than 1.5 meters. For this study, sampling was carried out in a single polygon with an area of 0.49 hectares located in the village of La Vuelta, Tibacuy.

**Coffee plantations:** these areas are characterized by developing in patches under shade, which could be temporary or permanent, generated by arboreal elements or with free exposure to sunlight. Six crops were selected, including five in La Vuelta (Tibacuy) and one in Buenos Aires (Nilo). The study considered crop maintenance variables, such as the use of agrochemicals, periodicity of irrigation and maintenance, as well as the type of crop, which could be with or without shade, monoculture, or mosaic. In addition, four age classes were established for the crops according to the average plant height: Stage 1 (plants with heights less than 50 cm), Stage 2 (heights between 50 and 100 cm), Stage 3 (heights between 100 and 150 cm) and Stage 4 (heights greater than 150 cm). The abundance and humidity of leaf litter were recorded, as well as the presence of rocks, trunks, and other potential microhabitats.

**Urban areas:** these are composed of buildings and surrounding green spaces. In the study area, a private property located in the Buenos Aires (Nilo) neighborhood with an area of 0.068 ha was included. This site is surrounded by a rural road, gardens, and green spaces. Although urban areas were not part of the initial experimental design due to its anthropized nature with low diversity potential, they were included due to the incidental observation of *Hemidactylus* sp. during logistical activities associated with the field stage.

**Table 2.**
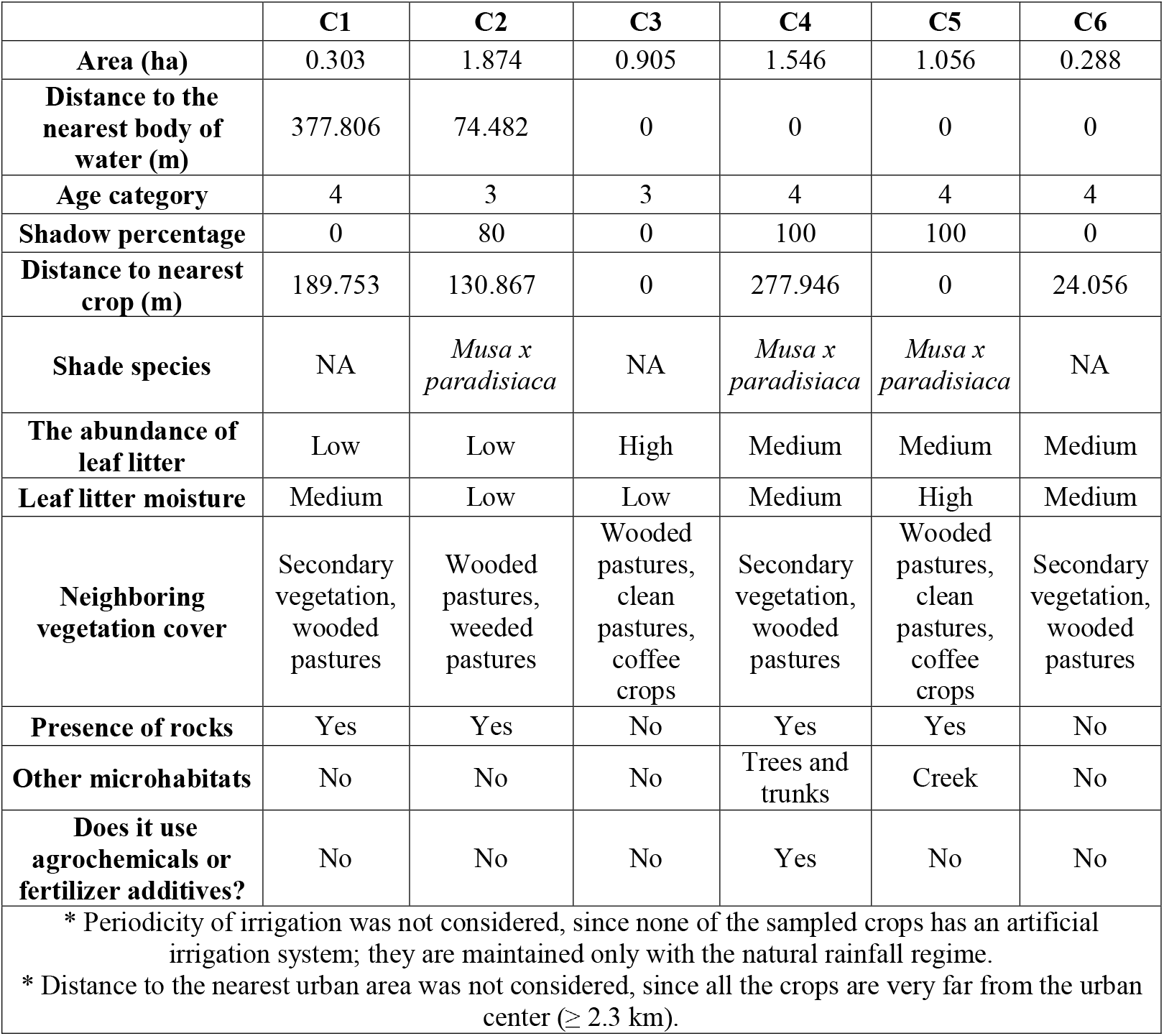
Structural and maintenance features of coffee crops. This table presents the characteristics of the six sampled coffee crops. None of the sampled crops uses agrochemicals or fertilizer additives, and the periodicity of irrigation was not considered as all crops rely only on the natural rainfall regime. Distance to the nearest urban area was also not considered, as all crops were located at least 2.3 km away from the urban center.

|  | C1 | C2 | C3 | C4 | C5 | C6 |
| --- | --- | --- | --- | --- | --- | --- |
| **Area (ha)** | 0.303 | 1.874 | 0.905 | 1.546 | 1.056 | 0.288 |
| **Distance to the nearest body of water (m)** | 377.806 | 74.482 | 0 | 0 | 0 | 0 |
| **Age category** | 4 | 3 | 3 | 4 | 4 | 4 |
| **Shadow percentage** | 0 | 80 | 0 | 100 | 100 | 0 |
| **Distance to nearest crop (m)** | 189.753 | 130.867 | 0 | 277.946 | 0 | 24.056 |
| **Shade species** | NA | *Musa x paradisiaca* | NA | *Musa x paradisiaca* | *Musa x paradisiaca* | NA |
| **The abundance of leaf litter** | Low | Low | High | Medium | Medium | Medium |
| **Leaf litter moisture** | Medium | Low | Low | Medium | High | Medium |
| **Neighboring vegetation cover** | Secondary vegetation, wooded pastures | Wooded pastures, weeded pastures | Wooded pastures, clean pastures, coffee crops | Secondary vegetation, wooded pastures | Wooded pastures, clean pastures, coffee crops | Secondary vegetation, wooded pastures |
| **Presence of rocks** | Yes | Yes | No | Yes | Yes | No |
| **Other microhabitats** | No | No | No | Trees and trunks | Creek | No |
| **Does it use agrochemicals or fertilizer additives?** | No | No | No | Yes | No | No |

\* Periodicity of irrigation was not considered, since none of the sampled crops has an artificial irrigation system; they are maintained only with the natural rainfall regime.
\* Distance to the nearest urban area was not considered, since all the crops are very far from the urban center ( $\geq 2.3$ km).

During the sampling phase, the protocol of Villarreal et al. (2006) was followed using the techniques of visual and auditory encounter survey (VES) for a limited time and fixed band transects (2 x 50 m). Sampling was conducted between 8:00 and 12:00 and between 18:00 and 22:00 hours, and potential habitats were searched for species, including vegetation, edges of ponds, streams, roads, and leaf litter, with a maximum limit of 5 meters above the ground (Cortés-Gómez et al. 2008).

In addition to the field trips conducted in 2021, historical records from field days between 2015 and 2019 were also included in the diversity analyses. These records were collected by the authors of this study and the herpetology group of Universidad Nacional de Colombia (Herpetos UN). During the field trips, data were collected on the site of observation, habitat, body measurements, and photographic records for all individuals. The individuals were released at the same location where they were captured. Furthermore, the cultivation systems were characterized based on proximity to water bodies, frequency of maintenance, use of agrochemicals, total area, adjacent vegetation cover, presence of microhabitats such as leaf litter and rocks, and percentage of shading. The software QGIS 3.22 (QGIS Development Team, 2023) was used for the spatial analysis.

### Data analysis

To determine the representativeness of the sampling, the species accumulation curve was used as per the guidelines established by Villarreal et al. (2006). The data matrix was analyzed using CHAO 1, ACE, and Cole’s Rarefaction estimators, with data based on abundance. Alpha diversity was calculated using the Margalef (specific richness index), Simpson (dominance index), and Shannon-Wiener (evenness index) indices. The Jaccard similarity index was used for beta diversity. All diversity indices were calculated using R software (R Core Team, 2022).

To evaluate the relationship between structural characteristics associated with coffee crops and the diversity of herpetofauna found in the crops, a canonical correspondence analysis (CCA) was performed using the CCA package in R software (Gonzalez et al., 2008; ter Braak, 1986), The structural characteristics included area, distance to the nearest water body, age category, shading, leaf litter abundance, leaf litter moisture, and distance to nearest crop. To ensure statistical robustness, the eigenvalue reliability criterion (Eigenvalue) ≥ 50 % was used.

## RESULTS

We captured a total of 134 individuals belonging to 33 species in the area: 14 species of frogs, 12 species of snakes, and 7 species of lizards (Supplementary material), turtles and crocodiles were not found in the sampling. This amounts to 48.3% of the potential amphibian species and 13.63% of the potential reptile species for the study area. The species accumulation curve (Figure 1) indicates that the samples taken in the RFPCQ are considerably representative, as it shows a tendency to reach the asymptote for the estimators used (ACE, CHAO 1, and Cole).

**Figure 1.**
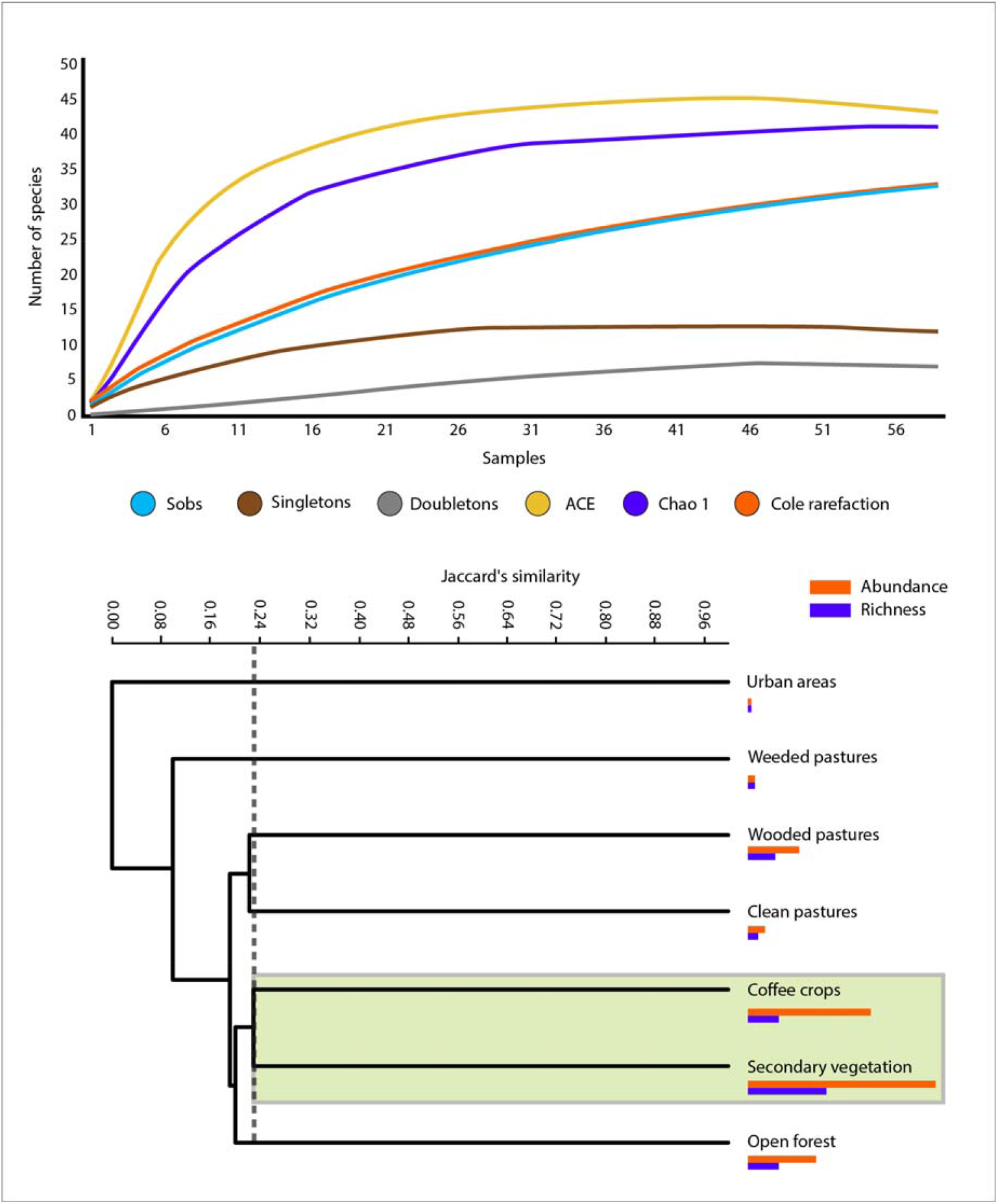
Sampling representativeness and species diversity across vegetation cover units. The curves depicted above approach the asymptote, indicating that the sampling is representative of the studied area. Representativeness percentage ranges from 77 to 100% (Villarreal et al., 2006). The Jaccard similarity index is presented for each vegetation cover unit to illustrate species diversity. The colored lines correspond to each unit’s absolute abundance (orange) and species richness (purple). The shaded area in the figure represents the clade with the highest Jaccard similarity value, which is approximately 25%. The dotted line indicates the cut-off point for the similarity of the most similar group.

The diversity of herpetofauna in the RFPCQ was analyzed by considering seven different cover units. Secondary vegetation had the highest number of recorded taxa and abundance, followed by coffee plantations and open forest. In contrast, the urban areas and weedy pasture units had the lowest values for both metrics (Table *1*). Simpson (1-D) and Shannon_H indices showed that secondary vegetation and wooded pastures were the most diverse coverages, while weeded pastures were the areas with the lowest diversity. On the other hand, the Margalef index revealed that both secondary vegetation and open forest were the most diverse, while cleared pastures had the lowest diversity (Table *1*).

In terms of beta diversity, Jaccard’s grouping index showed differences in the composition of species identified in distinct vegetation units. The urban areas and weeded pastures were highly different from the other units. Clean and wooded pastures were grouped on one hand, and coffee and secondary vegetation units on the other (Figure *1*). The latter showed considerable similarity with the composition of herpetofauna found in the open forest.

The canonical correspondence analysis (CCA) grouped 57.62% of the variation in the first two axis (Figure 2). The analysis revealed that the most important structural variables for determining herpetofauna diversity in the crop were leaf litter abundance, area, and age category. Litter moisture and distance to other crops also had a moderate influence on the presence of amphibians and reptiles. However, the distance to the nearest water body did not appear to be a good predictor of the data distribution.

**Figure 2.**
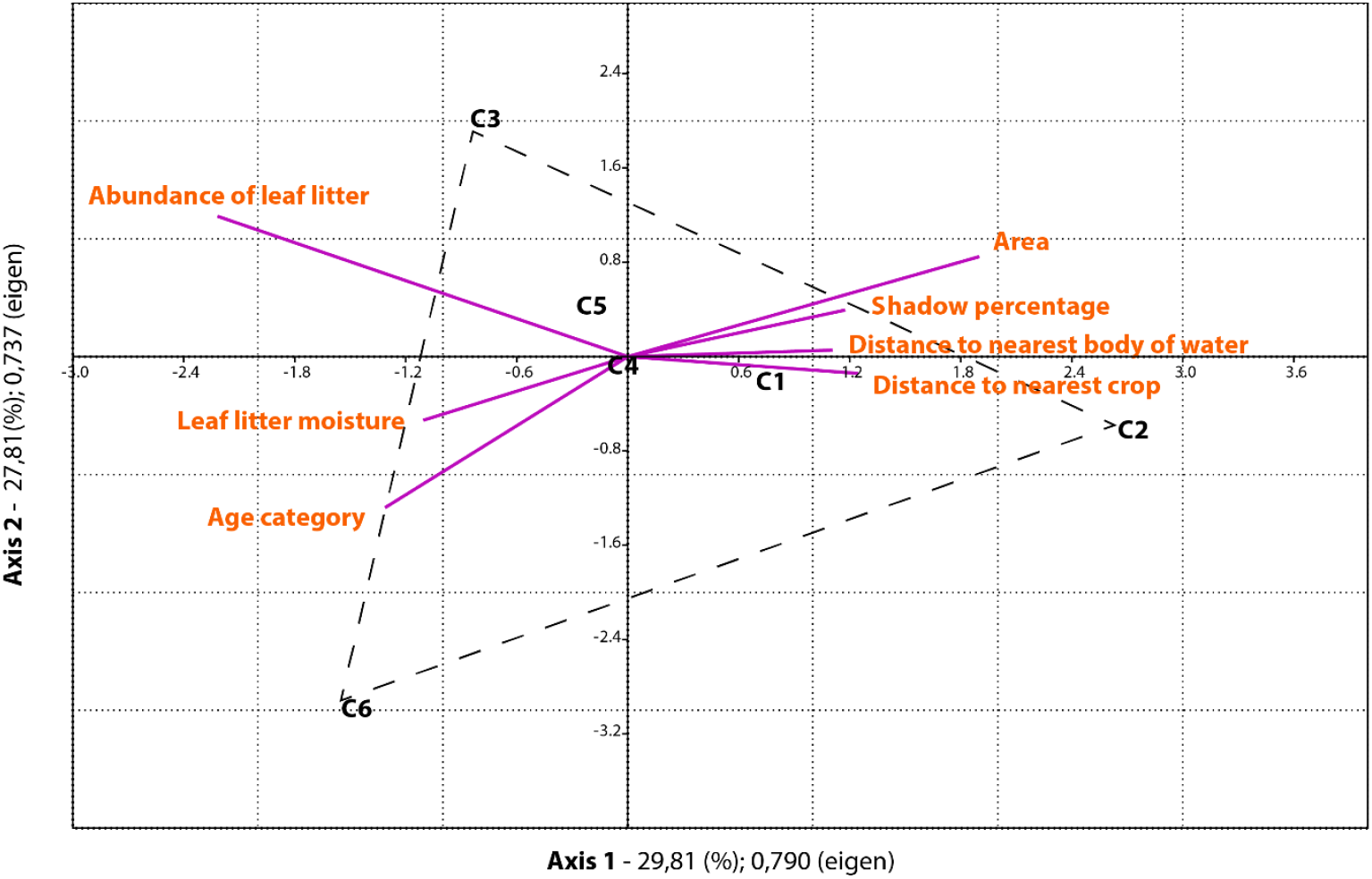
Canonical Correspondence Analysis (CCA). The diagram shows the species diversity relationship with the structural variables of 6 coffee crops in the RFPCQ. The length of the purple lines indicates the relative importance and direction of change of the structural variables. The title of each axis shows the cumulative percentage of variance explained (%) and the eigenvalue (eigen).

## DISCUSSION

The RFPCQ in the Sumapaz region has a diverse herpetofauna, with most species associated with secondary vegetation, forests, and coffee plantations characterized by shady conditions and advanced ages. These crops function similarly to forests and offer habitats conducive to the establishment of diverse groups of populations, allowing for greater diversity and non-generalist species to reside, even at considerable distances from the remaining forest (Rojas Sánchez et al., 2012; Tejeda-Cruz and Sutherland, 2004).

The analysis of the microclimatic variables showed the abundance of leaf litter, and the crop area has the greatest influence on diversity within the crops (Figure 2). Surprisingly, proximity to bodies of water, which is usually considered crucial for the diversity and abundance of amphibians, did not show to be a good predictor of the behavior of the data distribution. Studies have shown that the surrounding vegetation plays a significant role in maintaining relative humidity within a patch, resulting in better humidity conditions in coffee crops under shade than in crops under other types of management (Santos-Barrera and Urbina-Cardona, 2011).

Leaf litter is an essential microhabitat for herpetofauna, providing a moist environment even in the dry season (Cortés-Gómez et al., 2008). A thick layer of leaf litter within the crops could favor the establishment of complex and structured animal communities (Rojas Sánchez et al., 2012; Tejeda-Cruz and Sutherland, 2004), increasing the levels of diversity observed in agricultural environments. Furthermore, recent studies have shown that practices related to intensive coffee cultivation can promote population decline in amphibians due to the decrease in habitat quality and quantity of microhabitats. For example, in the Sierra Nevada de Santa Marta, the fourth-largest coffee-growing region in the country, population decline has been observed due to the expansion of the agricultural frontier (Roach et al., 2021, 2020). However, if coffee cultivation practices under shade in the RFPCQ are continued with the promotion of the maintenance of a thick layer of leaf litter in the coffee plantations and practices used so far, such as the non-use of fertilizers and pesticides, the diversity of amphibians and reptiles will not decline due to the persistence of these agroecosystems. This approach could also provide benefits to local communities by promoting environmentally friendly and socially responsible coffee production practices.

### A SET OF SUSTAINABLE MANAGEMENT PRINCIPLES TO PROMOTE HERPETOFAUNA COEXISTENCE

The findings of our study are in line with previous research indicating that shaded coffee plantations can provide suitable habitats for herpetofauna, including amphibians and reptiles. However, it is important to note that the use of agrochemicals, such as pesticides and fertilizers (not addressed in this study due to the specific area conditions), can have negative impacts on these animals and their habitats (Wagner et al., 2013). A recent study by Wagner et al. (2017) found that the use of glyphosate-based herbicides could induce morphological changes and increase malformation rates in amphibians, especially in areas with low forest cover. Therefore, it is crucial to adopt sustainable agricultural practices that minimize the use of agrochemicals and promote the maintenance of natural habitats.

Several principles and practices can be implemented to promote sustainable agricultural management of coffee crops that support the existence of amphibians, reptiles, and other terrestrial animals in the Andes. We propose the following 6 principles of management:

- Shade management: Maintaining a diverse shade cover that mimics the natural forest structure can provide suitable habitats for herpetofauna and other animals, as well as promote soil health and prevent erosion (Tscharntke et al., 2011).
- Leaf litter management: Maintaining a thick layer of leaf litter can provide a moist and structured environment that supports a diverse community of animals (Tejeda-Cruz and Sutherland, 2004), even in the dry season, which is crucial for their survival. However, it is important to avoid excessive accumulation of leaf litter, as this can lead to an increased risk of fire and pest outbreaks (Balch et al., 2015).
- Integrated pest management: Adopting pest control strategies that prioritize nonchemical methods, such as crop rotation, intercropping, and biological control, can reduce the negative impacts of agrochemicals on the environment and promote the natural regulation of pest populations (Crowder and Jabbour, 2014).
- Soil and water management: Practices such as the use of cover crops, composting, and reduced tillage can improve soil health and promote the retention of moisture and nutrients, which can benefit plant and animal communities (Bach et al., 2020). Also, implement sustainable water management practices, such as rainwater harvesting and irrigation systems that minimize water use and runoff, and avoid contamination of nearby water sources.
- Connectivity management: Promote landscape connectivity by preserving and restoring natural areas around coffee plantations, such as forests, wetlands, and other important habitats for terrestrial animals. These practices improve resource availability and favors the coexistence of plant and animal communities (Howell et al., 2018).
- Monitoring management: Implement monitoring and management plans to ensure that the above practices are being followed and that the health of the ecosystem is being maintained. This can include regular surveys of the herpetofauna and other wildlife and soil and water quality testing.

By implementing these sustainable practices, coffee farmers can contribute to the conservation of biodiversity and the maintenance of ecosystem services in coffee-growing regions, while also promoting the economic viability of their crops.

Furthermore, the promotion of agricultural practices that maintain the quality and quantity of microhabitats is crucial. This can be achieved by maintaining diverse successional stages within the coffee plantation, including secondary vegetation and forest. The presence of these habitats provides suitable conditions for the establishment of non-generalist species, even at considerable distances from the remaining forest. Moreover, maintaining the structural variables of leaf litter abundance, area, and age category as well as the moderate influence of litter moisture and distance to other crops will ensure the presence of amphibians and reptiles. Therefore, promoting sustainable agricultural management practices in coffee plantations can contribute to the conservation of the herpetofauna species and the overall biodiversity of the region.

In conclusion, the results of this study highlight the importance of agroecosystems such as shade coffee plantations in providing habitats for herpetofauna in highly fragmented landscapes such as the Sumapaz region. The diversity and abundance of amphibians and reptiles were found to be influenced by several microclimatic variables, with leaf litter abundance and crop area being the most important. The findings suggest that the maintenance of shade coffee plantations and the preservation of leaf litter layers within them can contribute to the conservation of herpetofauna diversity in highly modified landscapes.

Overall, the study contributes to the understanding of the ecological importance of agroecosystems in highly fragmented landscapes and the factors that influence the diversity and abundance of herpetofauna in these habitats. The findings have important implications for conservation strategies in the Sumapaz region and other areas with similar characteristics. Future research could focus on further understanding the dynamics of herpetofauna populations in agroecosystems and the interactions between agricultural practices, land use change, and biodiversity conservation in these environments.

## ACKNOWLEDGMENTS

We thank the Asociación Colombiana de Herpetología for funding through the Botas al Campo grant (2019) and APRENAT for their logistical support. We also acknowledge the cooperation of various individuals and organizations, including the Castillo Urrego family, Bernal Méndez family, Maria Rosa Gaona de Chacón, Mayelly Liévano, Gabriel Garzón, Fray José María Sepúlveda, the Province of Nuestra Señora de Gracia de Colombia and the members of Herpetos UN, for their hospitality, collaboration throughout the field phase and for their valuable contributions in the development of this project. To Nataly Casas, Carolina Martínez, Juan Diego Rodríguez, and Sebastián Pérez for their help during the field phase and species identification. Thanks to Kelley Crites for the English writing advising. Finally, we appreciate the ongoing efforts of the communities of La Vuelta, Capotes, El Cairo, Albania, and Buenos Aires in developing and conserving the RFPCQ.

## Funding

This work was supported by the Asociación Colombiana de Herpetología (ACH) through the Botas al Campo grant, 2019.

## Notes

https://figshare.com/articles/dataset/Supplementary_material_Conserving_Biodiversity_in_Coffee_Agroecosystems_Insights_from_a_Herpetofauna_Study_in_the_Colombian_Andes_with_Sustainable_Management_Proposal/22223761

### Competing Interest Statement

The authors have declared no competing interest.

