## Supplementary materials for "Conserving Biodiversity in Coffee Agroecosystems: Insights from a Herpetofauna Study in the Colombian Andes with Sustainable Management Proposal"

^3^ Independent researcher.

^4^ Grupo de Investigación de Ornitología, Instituto de Ciencias Naturales, Universidad Nacional de Colombia, Sede Bogotá, Colombia.

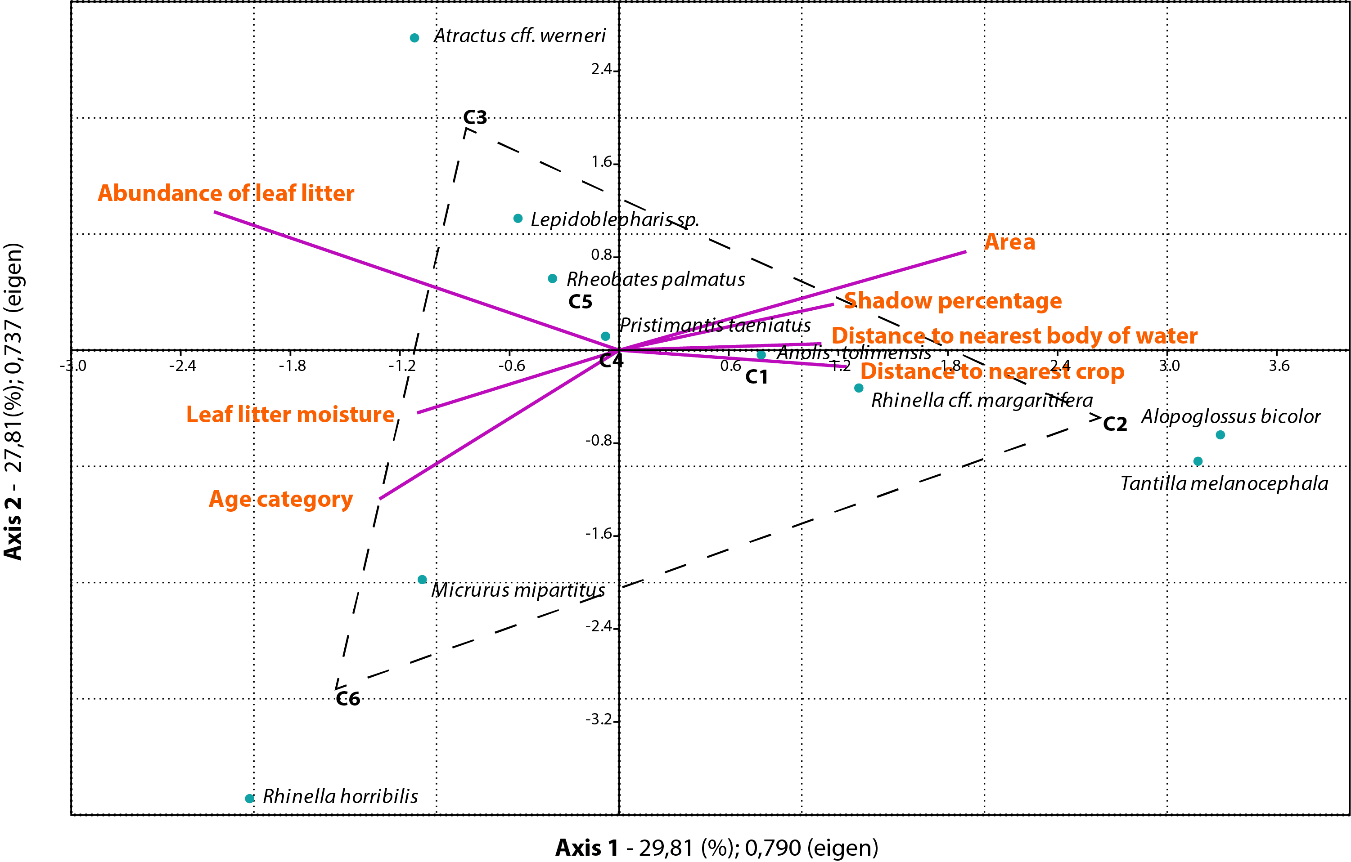

**Supplementary figure 1.** Canonical Correspondence Analysis (CCA) including species component distribution.

**Supplementary table 1.** Species of amphibians and reptiles identified in the study area. Of: open forest, Sv: secondary vegetation, Wp: wooden pastures, Wp: weeded pastures, Cp: clean pastures, Cc: coffee crops, Ua: urban areas.

| **Class** | **Order** | **Family** | **Genus** | **Species** | **Abundance** | **Of** | **Sv** | **Wp** | **Wep** | **Cp** | **Cc** | **Ua** |
| --- | --- | --- | --- | --- | --- | --- | --- | --- | --- | --- | --- | --- |
| Amphibia | Anura | Aromobatidae | Rheobates | *Rheobates palmatus* | 2 |  | 1 |  |  |  | 1 |  |
| Amphibia | Anura | Bufonidae | Rhinella | *Rhinella cf. margaritifera* | 5 |  | 2 |  |  |  | 3 |  |
| Amphibia | Anura | Bufonidae | Rhinella | *Rhinella horribilis* | 2 |  |  |  |  | 1 | 1 |  |
| Amphibia | Anura | Centrolenidae | Centrolene | *Centrolene daidalea* | 2 | 2 |  |  |  |  |  |  |
| Amphibia | Anura | Centrolenidae | Espadarana | *Espadarana andina* | 2 | 2 |  |  |  |  |  |  |
| Amphibia | Anura | Dendrobatidae | Dendrobates | *Dendrobates truncatus* | 3 |  | 1 | 2 |  |  |  |  |
| Amphibia | Anura | Hylidae | Boana | *Boana platanera* | 15 |  | 8 | 4 |  | 3 |  |  |
| Amphibia | Anura | Hylidae | Scinax | *Scinax gr. ruber* | 1 |  | 1 |  |  |  |  |  |
| Amphibia | Anura | Hylidae | Dendropsophus | *Dendropsophus padreluna* | 1 |  | 1 |  |  |  |  |  |
| Amphibia | Anura | Leptodactylidae | Leptodactylus | *Leptodactylus colombiensis* | 3 |  | 1 | 2 |  |  |  |  |
| Amphibia | Anura | Leptodactylidae | Leptodactylus | *Leptodactylus sp.* | 1 |  | 1 |  |  |  |  |  |
| Amphibia | Anura | Strabomantidae | Pristimantis | *Pristimantis gaigei* | 3 |  | 3 |  |  |  |  |  |
| Amphibia | Anura | Strabomantidae | Pristimantis | *Pristimantis savagei* | 1 | 1 |  |  |  |  |  |  |
| Amphibia | Anura | Strabomantidae | Pristimantis | *Pristimantis taeniatus* | 40 | 9 | 9 |  |  |  | 22 |  |
| Reptilia | Squamata | Boidae | Boa | *Boa constrictor* | 3 | 1 | 2 |  |  |  |  |  |
| Reptilia | Squamata | Colubridae | Atractus | *Atractus cf. werneri* | 2 |  |  | 1 |  | 1 |  |  |
| Reptilia | Squamata | Colubridae | Clelia | *Clelia sp.* | 1 |  | 1 |  |  |  |  |  |
| Reptilia | Squamata | Colubridae | Lampropeltis | *Lampropeltis micropholis* | 1 |  |  | 1 |  |  |  |  |
| Reptilia | Squamata | Colubridae | Leptodeira | *Leptodeira septentrionalis* | 1 |  | 1 |  |  |  |  |  |
| Reptilia | Squamata | Colubridae | Phrynonax | *Phrynonax poecilonotus* | 1 |  | 1 |  |  |  |  |  |
| Reptilia | Squamata | Colubridae | Spilotes | *Spilotes pullatus* | 1 |  | 1 |  |  |  |  |  |
| Reptilia | Squamata | Colubridae | Tantilla | *Tantilla melanocephala* | 2 |  |  |  | 1 |  | 1 |  |
| Reptilia | Squamata | Dactyloidae | Anolis | *Anolis apollinaris* | 3 | 1 | 2 |  |  |  |  |  |
| Reptilia | Squamata | Dactyloidae | Anolis | *Anolis tolimensis* | 11 |  | 7 | 3 |  |  | 1 |  |
| Reptilia | Squamata | Elapidae | Micrurus | *Micrurus dumerilii* | 2 |  | 1 |  | 1 |  |  |  |
| Reptilia | Squamata | Elapidae | Micrurus | *Micrurus mipartitus* | 3 | 1 |  |  |  |  | 2 |  |
| Reptilia | Squamata | Alopoglossidae | Alopoglossus | *Alopoglossus bicolor* | 5 | 1 | 3 |  |  |  | 1 |  |
| Reptilia | Squamata | Gekkonidae | Hemidactylus | *Hemidactylus sp.* | 1 |  |  |  |  |  |  | 1 |
| Reptilia | Squamata | Phyllodactylidae | Thecadactylus | *Thecadactylus rapicauda* | 3 |  | 2 | 1 |  |  |  |  |
| Reptilia | Squamata | Scincidae | Mabuya | *Mabuya sp.* | 3 | 2 | 1 |  |  |  |  |  |
| Reptilia | Squamata | Sphaerodactylidae | Lepidoblepharis | *Lepidoblepharis sp.* | 8 |  | 4 |  |  |  | 4 |  |
| Reptilia | Squamata | Viperidae | Bothrops | *Bothrops asper* | 1 |  | 1 |  |  |  |  |  |
| Reptilia | Squamata | Viperidae | Crotallus | *Crotallus durissus* | 1 |  |  | 1 |  |  |  |  |
| **Total** | | | | | 134 | 20 | 55 | 15 | 2 | 5 | 36 | 1 |
|  |  |  |  |  |  | 9 | 23 | 8 | 2 | 3 | 9 | 1 |

**Supplementary table 2.** Eigen values calculated for Canonical Correspondence Analysis (CCA).

| Eje | Eigenvalue | % |
| --- | --- | --- |
| 1 | 0.79043 | **29.81** |
| 2 | 0.73742 | **27.81** |
| 3 | 0.56233 | 21.21 |
| 4 | 0.3704 | 13.97 |
| 5 | 0.19058 | 7.189 |
